# β-Catenin Limits Osteogenesis on Regenerative Materials in a Stiffness-Dependent Manner

**DOI:** 10.1101/2021.06.15.447161

**Authors:** Qi Zhou, Xiaoyan Ren, Michelle K. Oberoi, Rachel M. Caprini, Marley J. Dewey, Vasiliki Kolliopoulos, Dean T. Yamaguchi, Brendan A.C. Harley, Justine C. Lee

**Affiliations:** Division of Plastic and Reconstructive Surgery, UCLA David Geffen School of Medicine, Los Angeles, CA 90095; Research Service, Greater Los Angeles VA Healthcare System, Los Angeles, CA 90073; UCLA Molecular Biology Institute, Los Angeles, CA 90095; Department of Chemical and Biomolecular Engineering, University of Illinois at Urbana-Champaign, Urbana, IL 61801; Carl R. Woese Institute for Genomic Biology, University of Illinois at Urbana-Champaign, Urbana, IL 61801

## Abstract

Targeted refinement of regenerative materials requires mechanistic understanding of cell-material interactions. The nanoparticulate mineralized collagen glycosaminoglycan (MC-GAG) scaffold is a porous biomaterial that promotes regenerative healing of calvaria defects *in vivo* without addition of exogenous growth factors or progenitor cells, suggesting its potential as an off-the-shelf implant for reconstructing skull defects. In this work, we evaluate the relationship between material stiffness, a tunable MC-GAG property, and activation of the canonical Wnt (cWnt) signaling pathway. Primary human bone marrow-derived mesenchymal stem cells (hMSCs) were differentiated on two MC-GAG scaffolds varying by stiffness (non-crosslinked, NX-MC, 0.3 kPa vs. conventionally crosslinked, MC, 3.9 kPa). hMSCs exhibited increased expression of activated β-catenin, the major cWnt intracellular mediator, and the mechanosensitive YAP protein with near complete subcellular colocalization in stiffer MC scaffolds. Small molecule Wnt pathway inhibitors reduced activated β-catenin and YAP protein quantities and colocalization, osteogenic differentiation, and mineralization on MC, with no effects on NX-MC. Concomitantly, Wnt inhibitors increased BMP4 and phosphorylated Smad1/5 (p-Smad1/5) expression on MC, but not NX-MC. Unlike non-specific Wnt pathway downregulation, isolated canonical Wnt inhibition with β-catenin knockdown increased osteogenic gene expression and mineralization specifically on the stiffer MC. β-catenin knockdown also increased p-Smad1/5, Runx2, and BMP4 expression only on the stiffer MC material. Our data indicates stiffness-induced activation of the Wnt and mechanotransduction pathways promotes osteogenesis in MC-GAG scaffolds. However, activated β-catenin is a limiting agent and may serve as a useful target or readout for optimal modulation of stiffness in skeletal regenerative materials.

**One Sentence Summary:** β-Catenin limits stiffness-induced osteogenenic differentiation on nanoparticulate mineralized collagen glycosaminoglycan materials

## Introduction

Materials-based approaches for tissue regeneration offer a unique potential to deliver simple, off-the-shelf products that are available at point of care in the operating room. In applications such as reconstruction of skull defects, the idea of cell-based or growth factor-based therapies may be impractical and, potentially, inferior when compared to the current clinical practices of autologous bone transfer or three-dimensionally printed alloplastic implants due to the need for additional invasive procedures including surgery for harvesting of progenitor cells, time required for progenitor cell expansion, or the risk of untoward consequences from supraphysiologic dosages of growth factors (*1*). However, one of the challenges in the development of biologically active implantable materials is that every composition, physical property, and modification thereof produces a unique cell biological response to the specific material of not just one cell type but all cell types within the microenvironment in which it is implanted (*2*). Hence, safe translation of biologically active implantable materials requires an improved understanding of cell biological responses induced by each individual material.

We previously described a nanoparticulate mineralized collagen glycosaminoglycan material (MC-GAG) that has shown promise as a cell-free, growth factor-free implantable material for skull regeneration in animal models (*3*). Hallmarks of this material include the ability to stimulate autogenous activation of the canonical bone morphogenetic protein receptor (BMPR) signaling pathway via phosphorylation of Smad1/5 (p-Smad1/5) in primary human mesenchymal stem cells (hMSCs) as well as primary rabbit bone marrow stromal cells (rBMSCs) (*3–5*). In addition, MC-GAG has both a direct and indirect inhibitory effect on primary human pre-osteoclasts (*6, 7*). While the trigger for such effects remains to be elucidated, we recently reported that stiffness of MC-GAG scaffolds resulted in activation of the mechanosensitive Yes-associated protein (YAP) and transcription activator with PDZ-binding motif (TAZ) signaling pathway with a concomitant increased expression of activated, non- phosphorylated β-catenin (non-p-β-catenin), the major intracellular mediator of the canonical Wingless-related integration site (cWnt) pathway (*8*). These data suggested that cWnt signaling may play an integral role in osteogenesis induced by MC-GAG in a stiffness-dependent manner.

The Wnt signaling pathways are known to be responsible for a multitude of functions including development, tissue homeostasis, and regeneration (*9*). In the cWnt pathway, Wnt ligands bind to one of the ten Frizzled (FZD) receptors with the low-density lipoprotein-related receptor 5 or 6 (LRP5 or LRP6) coreceptor thereby sequestering and inactivating the β-catenin destruction complex on the plasma membrane, liberating β-catenin for nuclear translocation (*10*). Interactions between YAP/TAZ and cWnt occur at the β-catenin destruction complex such that YAP and TAZ have both been co-immunoprecipitated with multiple proteins within the complex (*11, 12*). Nuclear β-catenin binds to the T-cell factor/lymphoid enhancer-binding factor (TCF/LEF) transcription factors and activates target genes (*9, 13*). Although significant crosstalk exists between the canonical and non-canonical Wnt pathways, the latter functions independently of β-catenin via two major pathways, the PCP (planar cell polarity)/JNK (Jun-n-terminal kinase) and calcium/PKC (protein kinase C) pathways (*14, 15*).

The effects of cWnt on osteogenic differentiation are varied. cWnt has been demonstrated to be necessary for osteoblast differentiation in murine cell lines and animal models (*16–18*). However, both stimulatory and inhibitory effects on osteogenic differentiation have been reported in human mesenchymal stem cells. Using small molecule inhibitors, Krause et al demonstrated that two different inhibitors which both increased β-catenin levels elicited contradictory effects on osteogenic differentiation of primary hMSCs (*19*). At least two groups have reported that activation of the canonical Wnt pathway inhibited osteogenic differentiation in primary hMSCs (*20, 21*). In the context of culturing on titanium surfaces, Olivares-Navarrete noted that non-canonical Wnt signaling was important for differentiation of osteoblast cell lines, whereas cWnt was inhibitory (*22*). These reports suggest that osteogenic or anti-osteogenic effects downstream of the Wnt pathways are likely dependent upon context including cell type, species, and environmental influences.

Within the realm of material development, the association between β-catenin activation in hMSCs and mechanical properties of MC-GAG scaffolds suggests that cWnt may serve as a useful molecular marker for tuning osteogenic activity in response to material stiffness. In this work, we evaluate the overall contributions of cWnt to the osteogenic activities of MC-GAG scaffolds as a function of material stiffness.

## Results

### Inhibition of the Wnt pathways preferentially reduce activation of β-catenin and mechanotransduction mediators on stiffer MC-GAG materials

We first evaluated the effects of the Wnt pathways using two small molecule inhibitors (*27*). IWR1 (inhibitors of Wnt response) targets the β-catenin destruction complex by stabilizing Axin, thereby potentiating β-catenin degradation (**Figure 1A**). IWP2 (inhibitors of Wnt production) targets Porcupine (Porcn), a member of the membrane-bound O-acyltransferase (MBOAT) family necessary for maturation of Wnt ligands via palmitoylation and secretion (*28*). While IWR1 primarily affects cWnt, IWP2 inhibits both canonical and non-canonical pathways.

**Figure 1.**
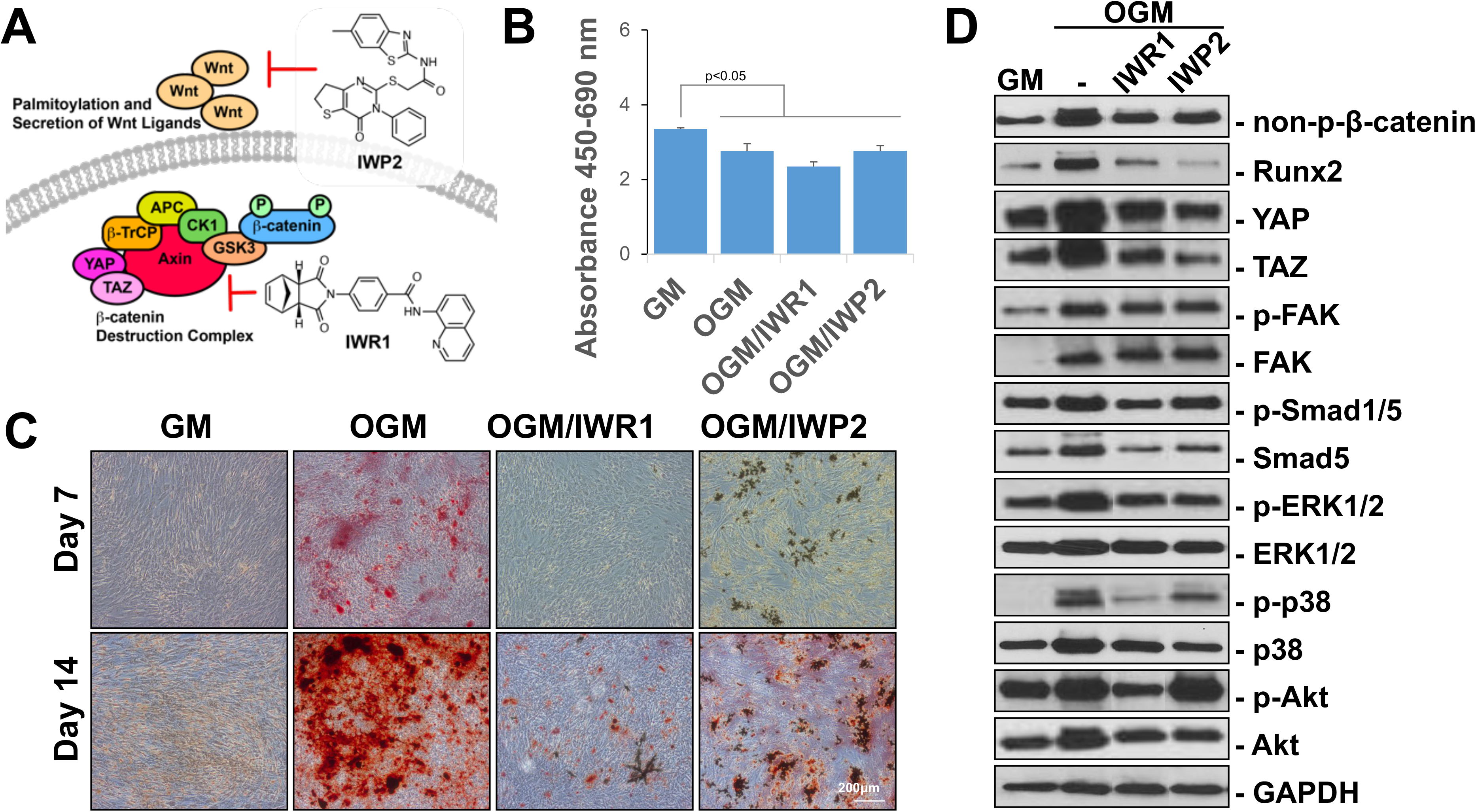
IWR1 and IWP2 inhibit accumulation of non-phosphorylated β-catenin and osteogenic differentiation of hMSCs in two-dimensional cultures. (**A**) Model of IWR1 and IWP2 mechanism of action. (**B**) WST-1 analysis of hMSCs cultured in GM, differentiated in OGM, or differentiated in OGM with 100 μM IWR1 (OGM/IWR1) or 50 μM IWP2 (OGM/IWP2) for 14 days. (C) Alizarin red staining of hMSCs cultured in GM, differentiated in OGM, or differentiated in OGM with 100 μM IWR1 (OGM/IWR1) or 50 μM IWP2 (OGM/IWP2) for 7 or 14 days. (**C**) Western blot of indicated intracellular mediators of hMSCs cultured in GM or OGM for 3 days in two-dimensional cultures. Cells were untreated or treated with 100 μM IWR1 or 50 μM IWP2. Bars represent means, errors bars represent SE. Significant posthoc comparisons following ANOVA indicated with p values.

To establish activity and dosage, primary bone marrow-derived hMSCs (CD105+CD166+CD29+CD44+CD14−CD34−CD45−) were cultured in growth media (GM) or differentiated in osteogenic growth media (OGM) in the absence and presence of each of the inhibitors in two-dimensional, monolayer cultures (**Figure 1B-D**).

Following a dose response analysis, the highest concentrations of IWR1 and IWP2 that resulted in both efficient downregulation of non-p-β-catenin without a significant reduction in housekeeping proteins (100 μM IWR1 and 50 μM IWP2) were chosen. To evaluate the effects of the inhibitors on cell viability and mineralization, hMSCs were cultured in GM or differentiated in OGM with or without each of the inhibitors for 14 days (**Figure 1B**). As expected, a reduction in proliferation was demonstrated when cells were cultured in osteogenic growth media versus basal growth media, likely corresponding to the increase in differentiation. Treatment with inhibitors did not change the viability or proliferation of cells. Mineralization of the cells were then assessed with Alizarin red staining at 7 days and 14 days of culture (**Figure 1C**). At both timepoints, both IWR1 and IWP2 reduced mineralization. To establish the baseline differences in signaling pathways, we also evaluated the impact of the two inhibitors on the expression of intracellular mediators in hMSCs induced to undergo osteogenic differentiation in monolayer, two-dimensional cultures (**Figure 1D**). Culture in OGM for 3 days demonstrated an increase in all intracellular mediators evaluated in comparison to culture in GM. IWR1 and IWP2 induced a reduction of non-p-β-catenin, Runx2, YAP, TAZ, p-ERK1/2, and p-p38. Effects on p-FAK, p-Akt, and p-Smad1/5 were more equivocal although a relative increase in p-Smad1/5 in relationship to total Smad5 was present.

After confirmation that both IWR1 and IWP2 were effective inhibitors of osteogenic differentiation and β-catenin activation in primary hMSCs in monolayer cultures, we then evaluated the inhibitors on NX-MC (0.34 ± 0.11 kPa) and MC (3.90 ± kPa) materials (**Figure 2A**). The two materials, as we previously described, have the exact same composition and are fabricated in an identical manner with the exception of increased stiffness in MC secondary to crosslinking performed after fabrication (*8*). Using WST-1 assays, we first determined whether either inhibitor affected viability and proliferation of hMSCs cultured on the two materials (**Figure 2B**). At 14 days of culture, no differences were found among hMSCs cultured without treatment, with IWR1 treatment, or with IWP2 treatment on either material.

**Figure 2.**
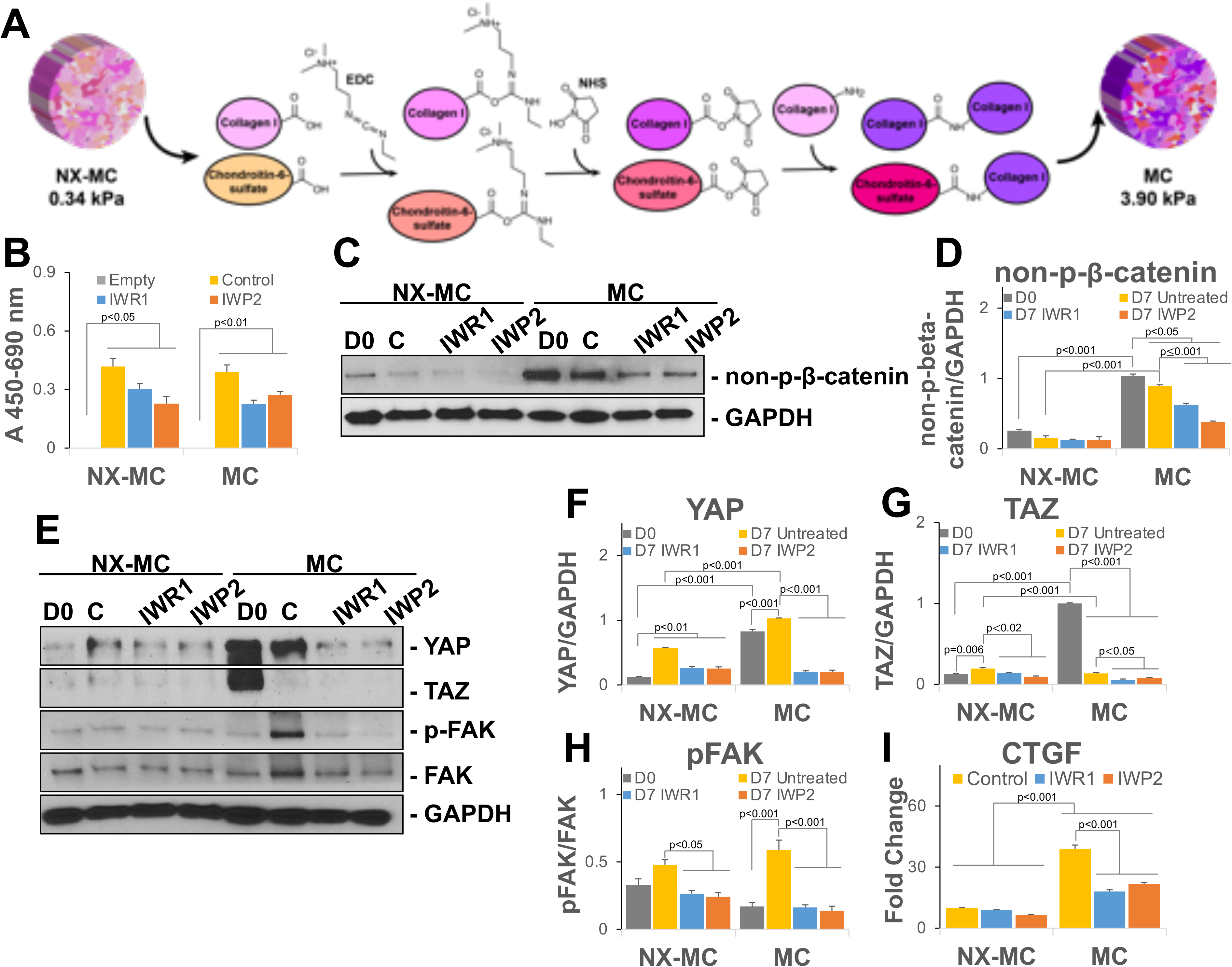
IWR1 and IWP2 downregulate non-phosphorylated β-catenin and mechanotransduction mediators preferentially on stiffer MC-GAG materials. (**A**) Schematic of NX-MC and MC mechanochemical differences with EDC/NHS crosslinking reaction depicted for generation of stiffer MC scaffold. (**B**) WST-1 analysis of NX-MC and MC cultured cell-free (Empty), with untreated hMSCs (Control), hMSCs treated with 50 μM IWR1 or IWP2 for 14 days. Representative image (**C, E**) or quantitative densitometric analyses of Western blots (**D, F-H)** for non-phosphorylated β-catenin (non-p-β-catenin), YAP, TAZ, phosphorylated FAK (p-FAK), FAK, and GAPDH in hMSCs differentiated on NX-MC and MC untreated (C), treated with IWR1, or treated with IWP2 for 7 days. Day 0 (D0) shown for comparison prior to differentiation. (**I**) QPCR of primary hMSCs cultured on NX-MC or MC for 7 days without (control) or with 50 μM IWR1 or IWP2 for CTGF. Bars represent means, errors bars represent SE. Significant posthoc comparisons following ANOVA indicated with p values.

Next, we evaluated the expression of non-p-β-catenin on the two materials in the absence and presence of IWR1 and IWP2 (**Figure 2C-D**). In the absence of inhibitors, the stiffer MC material induced significantly more non-p-β-catenin expression compared to NX-MC both prior to differentiation (day 0) as well as after 7 days of culture. Both qualitatively and quantitatively, IWR1 and IWP2 significantly reduced non-p-β-catenin expression when compared to untreated cultures specifically on the stiffer MC scaffold (p<0.001 for both), whereas no differences were seen on NX-MC suggesting that inhibition of cWnt or both canonical and non-canonical Wnt pathways preferentially affected activation of β-catenin on stiffer materials.

Due to the strong relationship between β-catenin activation and the mechanotransduction pathways, YAP, TAZ, and phosphorylated FAK (p-FAK) expression were also evaluated in the presence of IWR1 and IWP2 on the two materials (**Figure 2E-H**). On NX-MC, IWR1 and IWP2 treatment did not affect YAP expression when compared to untreated controls, however, both TAZ and p-FAK exhibited small reductions in relative expression in the presence of IWR1 and IWP2. hMSCs differentiated on MC differed from NX-MC in that YAP, TAZ, and p-FAK were expressed in greater quantities, albeit at different times during differentiation for the respective proteins. In the presence of IWR1 and IWP2, all three proteins displayed significant reductions in expression compared to untreated cultures. To understand whether the reduction in the mechanotransduction mediators translated to downstream reductions in transcriptional targets, we evaluated the expression of CTGF, a well-known target of YAP/TAZ activation (**Figure 2I**). As expected, CTGF expression was significantly higher in untreated hMSCs cultured on MC compared to NX-MC. While CTGF expression was unaffected by IWR1 and IWP2 on NX-MC materials, both inhibitors significantly reduced CTGF expression on MC scaffolds. In combination, these data suggest that inhibition of the canonical and/or non-canonical Wnt signaling pathways preferentially downregulated the activation of β-catenin on the stiffer MC material. Inhibition of the Wnt signaling pathways also simultaneously reduced YAP protein expression and expression of a YAP transcriptional target, CTGF, preferentially on the stiffer MC material.

### Inhibition of the Wnt pathways downregulate osteogenic gene expression and mineralization preferentially on stiffer MC-GAG materials

Next, we assessed the effects of Wnt inhibition on osteogenic differentiation and mineralization of hMSCs on NX-MC and MC (**Figure 3**). Primary hMSCs differentiated on NX-MC and MC for 7 days were evaluated for expression of osteogenic markers. For all osteogenic genes evaluated, hMSCs differentiated on MC demonstrated significantly higher expression compared to NX-MC (p<0.001 for all genes, **Figure 3A-D**). For expression of alkaline phosphatase (ALP), an early marker of differentiation, both IWR1 and IWP2 reduced expression on either NX-MC or MC materials. For collagen I (COL1A1), bone sialoprotein 2 (BSP2), and RUNX2, neither inhibitor significantly affected expression on NX-MC when compared to untreated controls. In contrast, IWR1 and IWP2 significantly reduced expression of all three former genes on stiffer MC materials. Of note, for ALP, COL1A1, BSP2, and RUNX2, gene expression was not completely eliminated by the inhibitors on stiffer MC materials. Rather, in the presence of the Wnt inhibitors, osteogenic gene expression was reduced to a level equivalent to hMSCs differentiated on the less stiff, NX-MC material.

**Figure 3.**
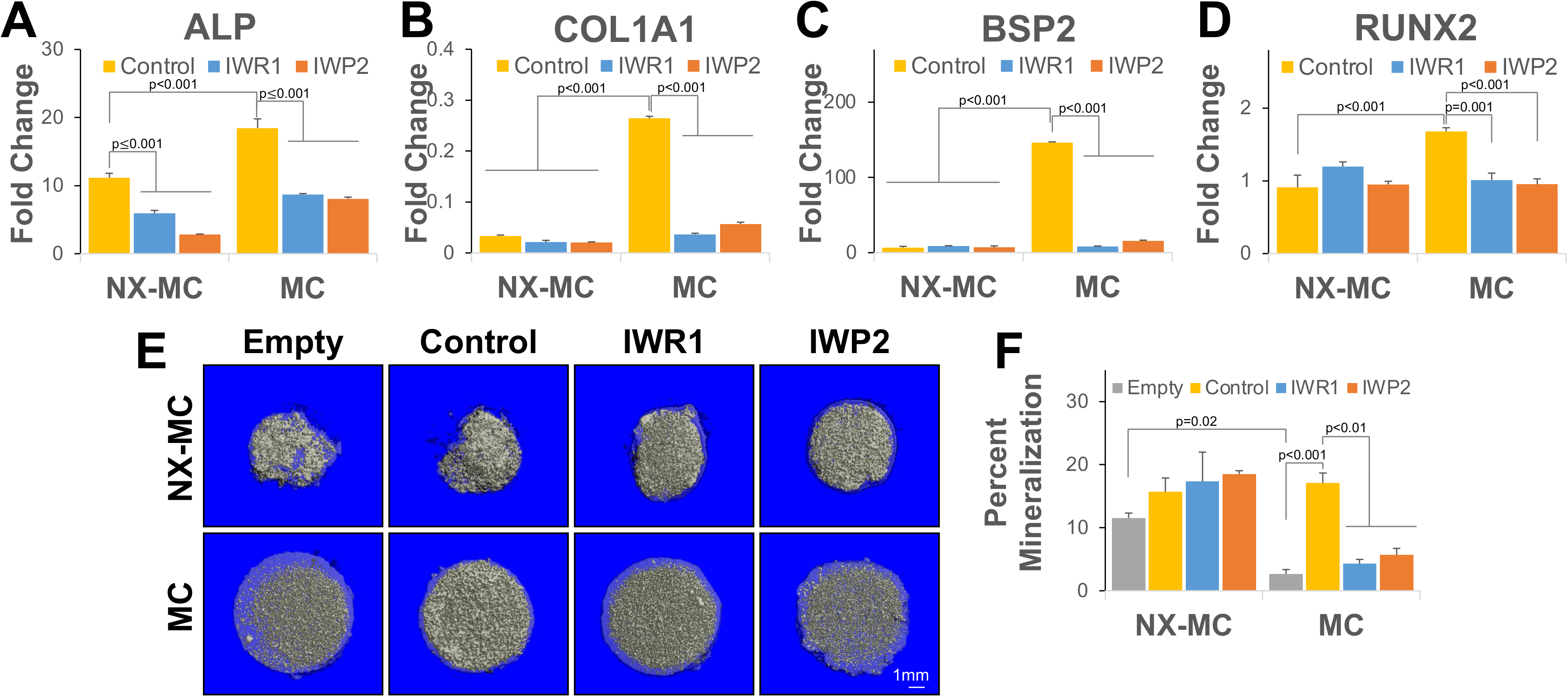
IWR1 and IWP2 downregulate osteogenic gene expression and mineralization preferentially on stiffer MC-GAG materials. QPCR of primary hMSCs cultured on NX-MC or MC for 7 days without (control) or with 50 μM IWR1 or IWP2 for (**A**) ALP, (**B**) COL1A1, (**C**) BSP2, and (**D**) RUNX2 (n=3). (**E**) Representative microCT images and (**F**) quantitative analysis NX-MC and MC cultured cell-free (Empty), with untreated hMSCs (Control), hMSCs treated with 50 μM IWR1 (IWR1) or 50 μM IWP2 (IWP2) for 8 weeks (n=4). Bars represent means, errors bars represent SE. Significant posthoc comparisons following ANOVA indicated with p values.

To understand if the differences in osteogenic gene expression were manifested in mineralization, hMSCs were cultured on NX-MC and MC in the absence and presence of IWR1 and IWP2 for 8 weeks and analyzed with microcomputed tomography (**Figure 3E-F**). Of note, the softer NX-MC scaffolds demonstrated significantly increased contraction, resulting in a smaller, denser material. Additional hMSC-mediated mineralization on NX-MC was relatively minimal compared to empty materials although the difference between pre-existing mineral on the scaffold versus cell-mediated deposition of mineral cannot be distinguished on microCT. Treatment with either IWR1 or IWP2 had no statistically significant effects on mineralization on NX-MC compared to untreated controls. In contrast, empty MC scaffolds were less dense compared to NX-MC and contraction during hMSC differentiation was relatively low. While a significant increase in mineralization occurred in stiffer MC scaffolds seeded with untreated hMSCs, treatment with either IWR1 or IWP2 severely reduced mineralization to levels similar to that of empty materials on MC (p<0.01). These data suggest that inhibition of the Wnt signaling pathways reduce osteogenic differentiation and mineralization of hMSCs specifically on stiffer MC-GAG materials.

### IWR1 and IWP2 induce a concomitant increase in BMP4 expression and Smad1/5 phosphorylation on stiffer MC-GAG materials

While the Wnt signaling pathways appear to play a role in osteogenic differentiation of stiffer MC materials, an obligate pathway for osteogenic differentiation on MC-GAG materials is the canonical BMPR signaling pathway via Smad1/5 activation (*3–5*). Hence, we next asked whether Smad1/5 and other intracellular mediators were differentially affected by Wnt inhibition on the two materials (**Figure 4**).

**Figure 4.**
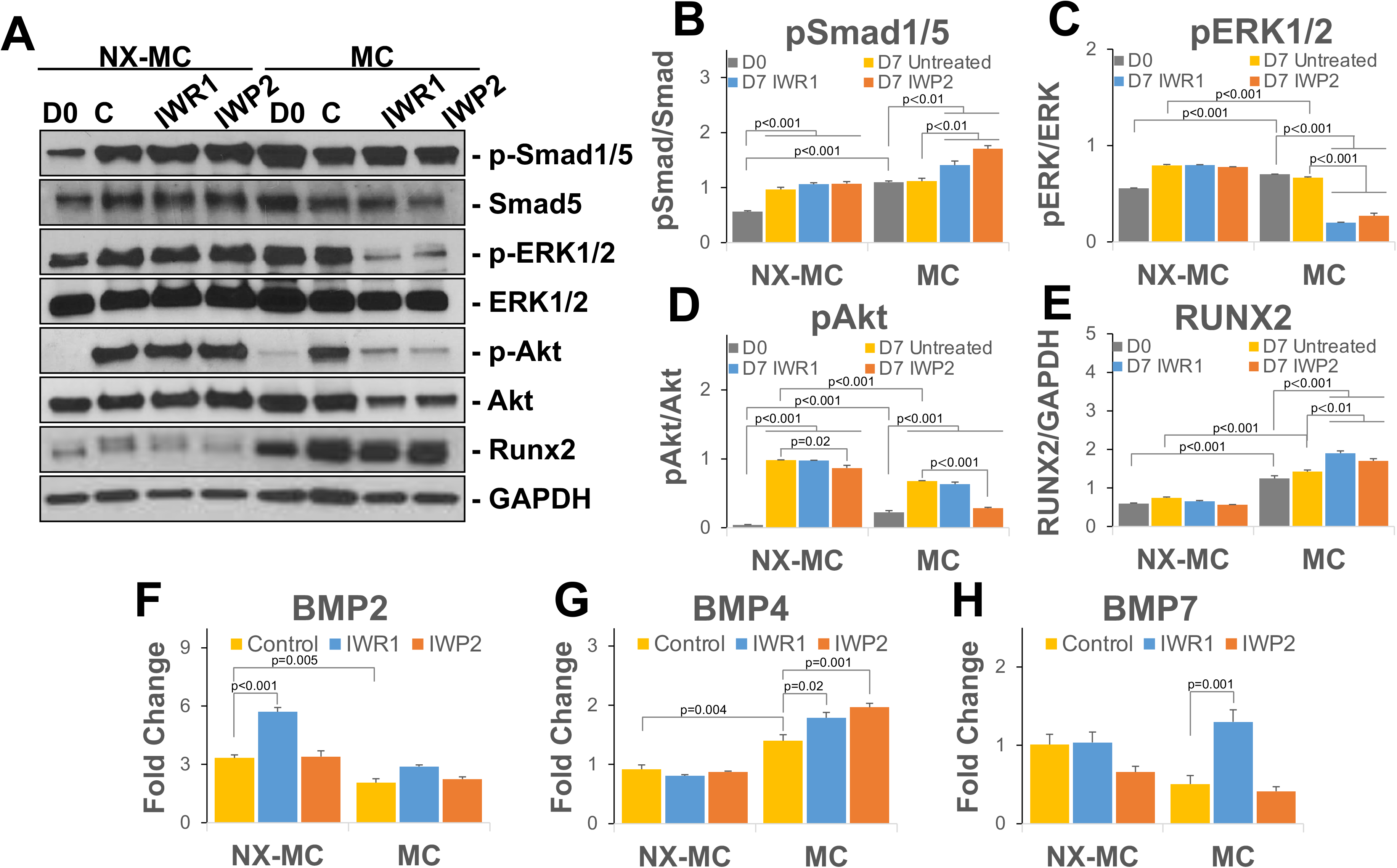
IWR1 and IWP2 induce a concomitant increase in BMP4 expression and Smad1/5 phosphorylation on stiffer MC-GAG materials. Representative image (**A**) or quantitative densitometric analyses of Western blots (**B-E)** for phosphorylated Smad1/5 (p-Smad1/5), total Smad5, phosphorylated ERK1/2 (p-ERK1/2), total ERK1/2, phosphorylated Akt (p-Akt), Akt, Runx2, and GAPDH. QPCR of primary hMSCs cultured on NX-MC or MC for 7 days without (control) or with 50 μM IWR1 or IWP2 for (**F**) BMP2, (**G**) BMP2, and (**H**) BMP7 (n=3). Bars represent means, errors bars represent SE. Significant posthoc comparisons following ANOVA indicated with p values.

In the presence of either NX-MC or MC materials, untreated control hMSCs demonstrated efficient activation of p-Smad1/5 with no statistically significant differences between the materials (**Figure 4A** and B). Inhibition with either IWR1 or IWP2 yielded no differences in the relative quantities of p-Smad1/5 to total Smad5 on NX-MC. In contrast, both inhibitors increased the quantity of p-Smad1/5 in relationship to total Smad5 when compared to the untreated scaffolds after 7 days of differentiation on the stiffer MC scaffold. A similar increase in quantitative Runx2 protein expression was also seen in the presence of IWR1 and IWP2 specifically on the stiffer MC material, yet absent in NX-MC scaffolds (**Figure 4E**). Unlike p-Smad1/5 and Runx2, both p- ERK1/2 and p-Akt displayed reductions on MC in the presence of IWR1 or IWP2 (**Figure 4A**,C, and D). For both p-ERK1/2 and p-Akt, minimal to no differences in expression were found in the presence of IWR1 or IWP2 on NX-MC compared to untreated controls.

While no exogenous BMP ligands were introduced into this system, the activation of the BMP receptor signaling pathway likely corresponded to an autogenous activation of BMP ligand expression. Thus, we also evaluated the expression of the dominant BMP ligands in hMSCs during osteogenic differentiation, BMP2, BMP4, and BMP7 (**Figure 4F-H**). BMP2 expression was consistently elevated in hMSCs cultured on NX- MC than on MC. In the presence of cWnt inhibition with IWR1, an increase in BMP2 expression was found in NX-MC but not stiffer MC scaffolds. IWP2-mediated inhibition had no effect on BMP2 expression on either material. By contrast, BMP7 expression showed no differences with or without Wnt inhibition on NX-MC, yet demonstrated a significant upregulation in the presence of IWR1 but not IWP2 on MC. BMP4 was also unaffected by either IWR1 or IWP2 on NX-MC, but was significantly upregulated by either inhibitor within stiffer MC scaffolds. In combination, these data suggest that inhibition of the canonical and/or noncanonical Wnt signaling pathway upregulates the BMPR signaling pathway via an increase in expression of BMP4 while downregulating other intracellular signaling mediators such as ERK1/2 and Akt in a stiffness-dependent manner. Additionally, inhibition of cWnt appears to upregulate BMP7 in a stiffness-dependent manner.

### Colocalization of activated β-catenin and YAP is reduced in the presence of Wnt inhibitors on stiffer MC-GAG materials

From both our gene and protein expression analyses, multiple lines of evidence suggested that the activity and protein quantities of non-p-β-catenin paralleled YAP on stiffer MC materials. Thus, we next evaluated whether the two proteins would colocalize in the same subcellular compartments (**Figure 5**).

**Figure 5.**
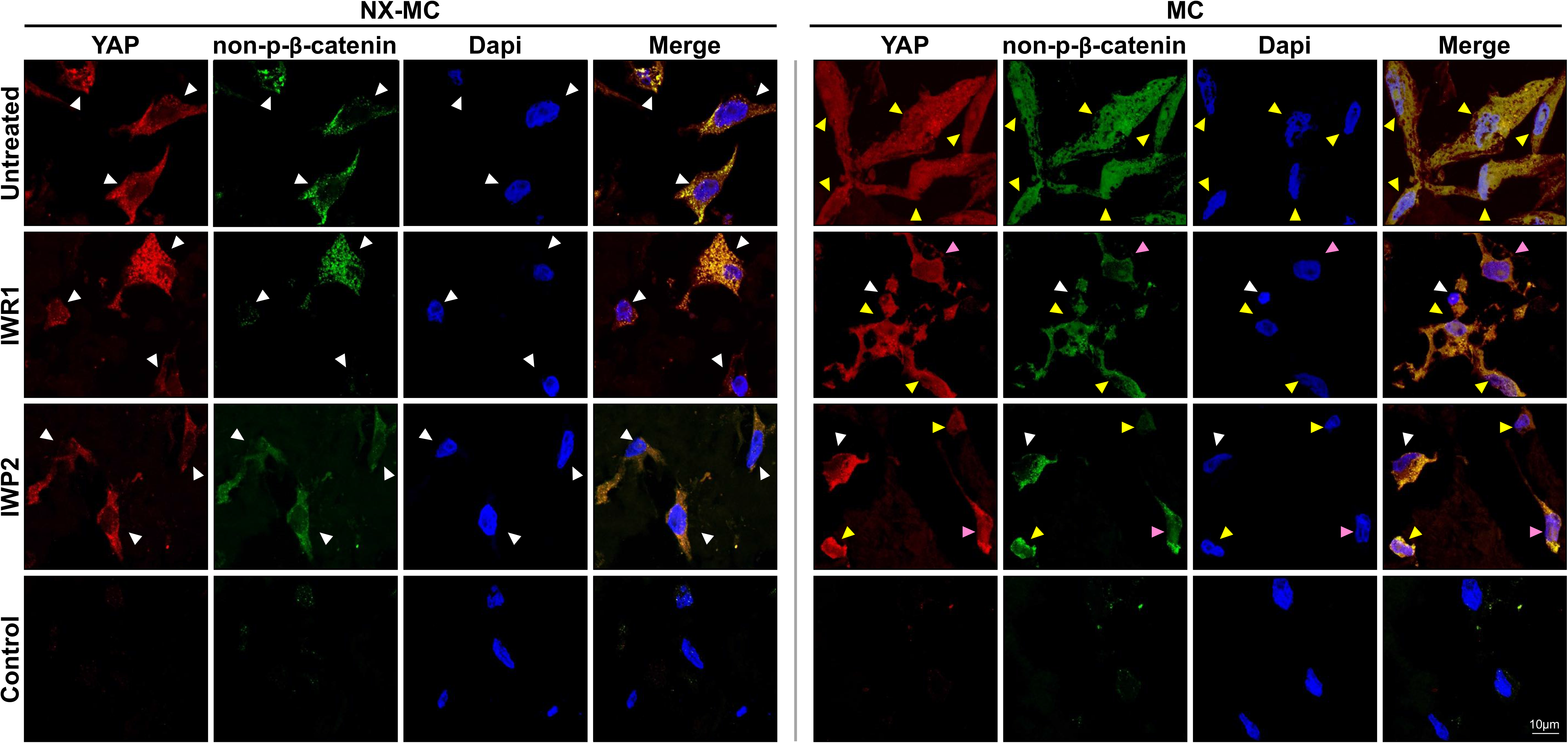
Colocalization of activated β-catenin and YAP is reduced in the presence of Wnt inhibitors on stiffer MC-GAG materials. Representative confocal microscopic images of primary hMSCs cultured on NX-MC (left panels) or MC (right panels) for 7 days untreated (top row) or treated with 50 μM IWR1 (second row) or IWP2 (third row) and stained for YAP or non-p-β-catenin. Negative control (Control) using secondary antibody only and Dapi on cells cultured on MC shown on bottom row. Scale bar indicates 10 μm. White arrows indicate cells with primarily cytosolic staining, yellow arrows indicate cells with both cytosolic and nuclear staining, and pink arrows indicate cells where YAP and non-p-β-catenin demonstrate a reduction in colocalization.

Primary hMSCs cultured on NX-MC for 7 days demonstrated primarily cytosolic localization of both YAP and non-p-β-catenin in the untreated controls as well as cells treated with IWR1 and IWP2. The increased stiffness in the MC material changed the subcellular localization of the two proteins such that both were found in the cytosol and the nucleus with nearly exact colocalization. In the presence of IWR1 and IWP2, a significant reduction in overall and nuclear staining of non-p-β-catenin occurred on MC scaffolds. While similar reductions were observed in YAP staining, a distinct reduction in colocalization of the two proteins was apparent. In a number of cells treated with IWR1 and IWP2, YAP staining continued to be relatively strong in the nucleus (pink arrows), whereas non-p-β-catenin was primarily found in the cytosol. These data suggest that Wnt inhibition also affects the YAP-mediated mechanotransduction pathway, however, YAP likely has functions independent of the Wnt pathway during osteogenic differentiation on stiffer MC materials.

### β-catenin knockdown regulates expression of mechanotransduction mediators and osteogenic gene expression in a stiffness dependent manner on MC-GAG materials

Despite providing a clear association between Wnt pathway inhibition and reduction in osteogenic differentiation on stiffer MC materials, two limitations exist with the usage of small molecule inhibitors: 1. The inability to separate the effects of YAP/TAZ mediated mechanotransduction and 2. The inability to separate the requirement of cWnt from the non-canonical Wnt pathways. Thus, to understand the role of cWnt pathway in isolation on osteogenic differentiation on the differential materials, we knocked down β-catenin expression using RNA interference against CTNNB1 (siCTNNB1) in primary hMSCs (**Figure 6**).

**Figure 6.**
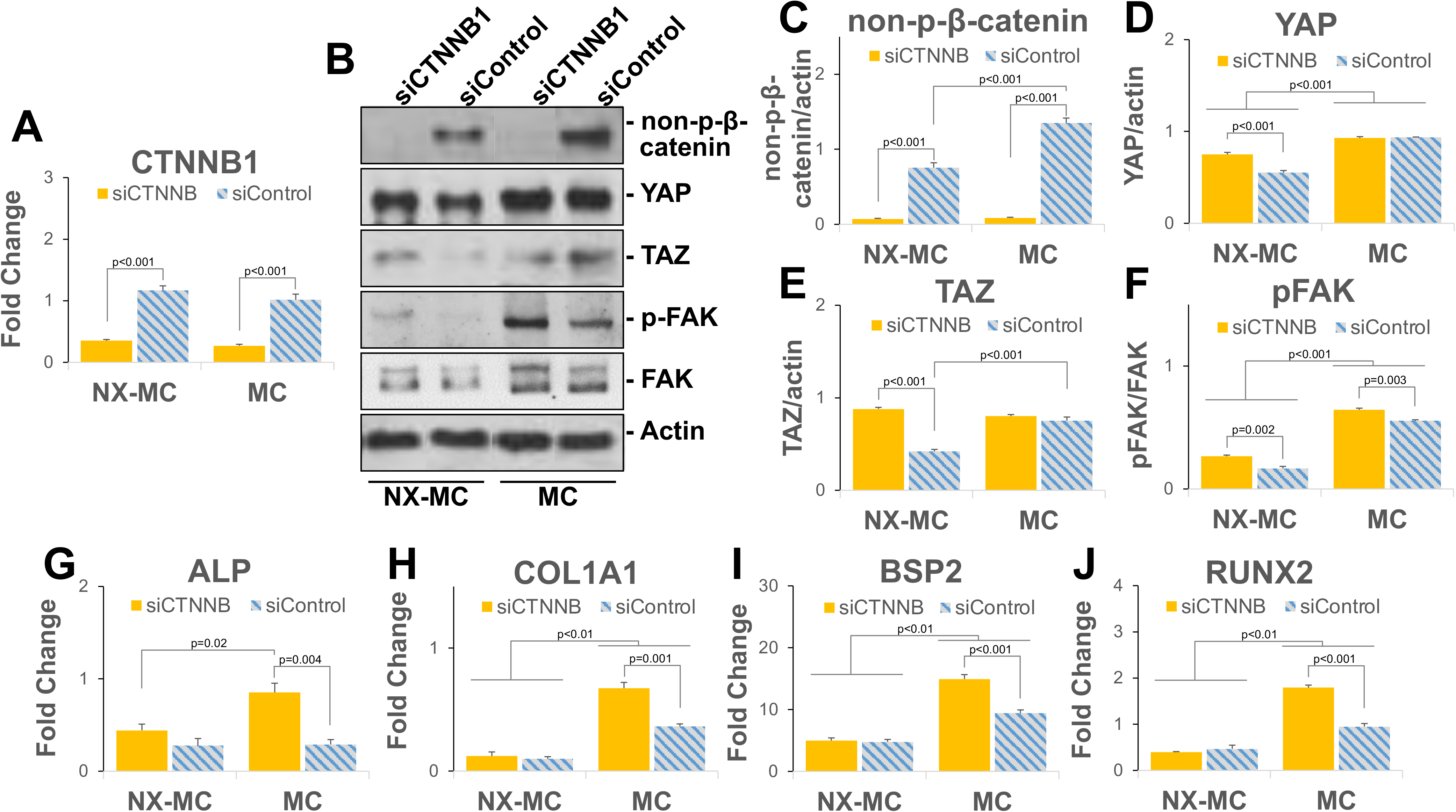
β-catenin knockdown regulates mechanotransduction mediators and osteogenic gene expression in a stiffness dependent manner. QPCR of primary hMSCs transfected with small interfering RNA targeting CTNNB1 (siCTNNB) or scrambled control (siControl) and cultured on NX-MC or MC for 7 days for (**A**) CTNNB1, (**G**) ALP, (**H**) COL1A1, (**I**) BSP2, and (**J**) RUNX2 (n=3). (**B**) Representative images and (**C-F**) quantification of Western blots of primary hMSCs transfected with siCTNNB or siControl and cultured on NX-MC or MC for 7 days for non-p-β-catenin, YAP, TAZ, p-FAK, FAK, and actin. Bars represent means, errors bars represent SE. Significant posthoc comparisons following ANOVA indicated with p values.

In the presence of siCTNNB1, a significant reduction in β-catenin gene expression as well as non-p-β-catenin protein expression was found in hMSCs cultured on either material when compared with cells transfected with scrambled control siRNA (siControl) (**Figure 6A-B**). Compared to inhibition of the Wnt pathway using IWR1 and IWP2, siCTNNB1 elicited distinct effects on YAP, TAZ, and p-FAK expression (**Figure 6C-F**). While the small molecule Wnt inhibitors generally reduced the expression of YAP, TAZ, and p-FAK, siCTNNB1 induced an overall increase in the expression of all three proteins on NX-MC materials. On the stiffer MC materials, siCTNNB1 did not induce any significant differences in YAP and TAZ expression while p-FAK was increased compared to siControl. These data suggest that overall inhibition of the Wnt pathways correlate to an inhibition of the mechanotransduction pathways, however, β- catenin appears to be a negative regulator of the mechanotransduction pathways such that knockdown of β-catenin actually increases YAP, TAZ, and p-FAK in softer MC-GAG materials with a lesser effect on the stiffer material.

We next evaluated the expression of osteogenic genes in the presence of siCTNNB1 on NX-MC and MC (**Figure 6G-J**). Expression of ALP, COL1A1, BSP2, and RUNX2 demonstrated similar expression patterns. All four genes were increased with siCTNNB1 on the stiffer MC material compared to siControl, whereas no differences were seen on NX-MC. Similar to the results in Figure 3, all four genes were expressed at higher levels on the stiffer MC material compared to NX-MC.

### β-catenin knockdown increases mineralization on stiffer MC-GAG materials with a concomitant increase in BMPR signaling

To understand if the upregulation in osteogenic gene expression on stiffer materials ultimately translated to differences in mineralization, hMSCs transfected with siCTNNB1 or siControl were differentiated on NX-MC or MC for 14 days or 8 weeks and subjected to Alizarin red staining or micro-computed tomographic imaging, respectively (**Figure 7A-D**). Similar to mineralization in the presence of the small molecule Wnt inhibitors, β-catenin knockdown had no effect on mineralization on softer NX-MC materials assessed by either Alizarin red staining or microCT. In the presence of MC materials, β-catenin knockdown significantly increased mineralization compared to control scaffold cultures. With either assay at either timepoint, the overall fold increase transfected cells.

**Figure 7.**
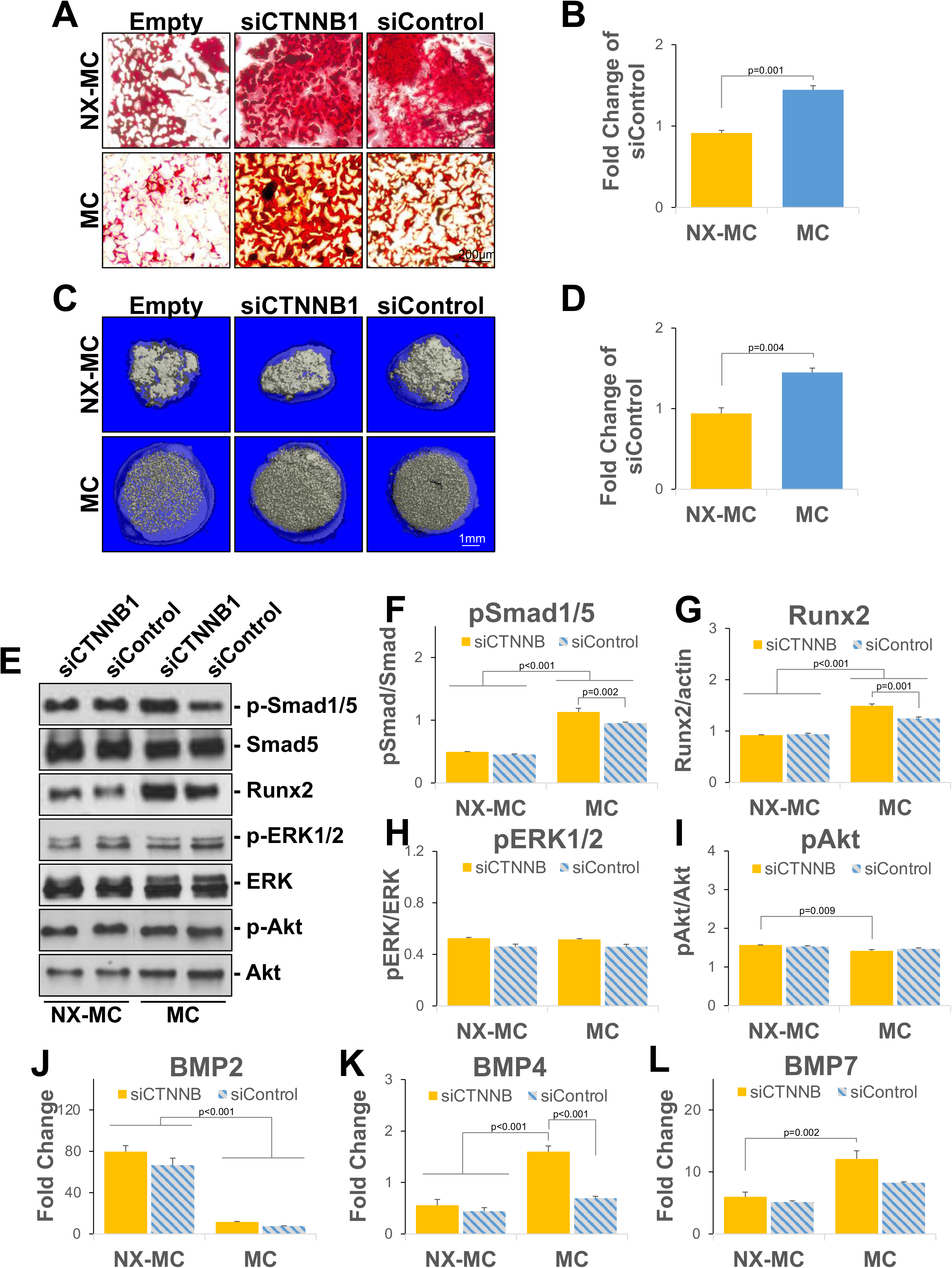
β-catenin knockdown increases mineralization on stiffer MC-GAG materials with a concomitant increase in BMPR signaling. (**A**) Representative images of Alizarin red staining of 4 micron sections of empty scaffolds (Empty) or hMSCs transfected with siCTNNB1 or siControl and differentiated on NX-MC or MC for 14 days (n=3). (**C**) Representative images of microCT of empty scaffolds (Empty) or hMSCs transfected with siCTNNB1 or siControl and differentiated on NX-MC or MC for 8 weeks (n=3). (**B, D**) Quantitative changes in mineralization in siCTNNB1 transfected cells on respective scaffolds expressed as percent mineralization in Alizarin red staining or microCT of siControl transfected cells. (**E**) Representative images and (**F-I**) quantification of Western blots of primary hMSCs transfected with siCTNNB or siControl and cultured on NX-MC or MC for 7 days for p- Smad1/5, Smad5, Runx2, p-ERK1/2, ERK1/2, p-Akt, and Akt. QPCR of primary hMSCs transfected with siCTNNB1 or siControl and cultured on NX-MC or MC for 7 days for (**J**) BMP2, (**K**) BMP4, and (**L**) BMP7 (n=3). Bars represent means, errors bars represent SE. Significant posthoc comparisons following ANOVA indicated with p values.

On evaluation of the BMPR signaling pathway, similar to IWR1 and IWP2 treatment, β-catenin knockdown increased p-Smad1/5 and Runx2 protein expression preferentially on stiffer MC materials whereas no differences were noted in the softer NX-MC material (**Figure 7E-G**). Minimal differences in phosphorylation of ERK1/2 or Akt were noted on either scaffold with β-catenin knockdown (**Figure 7H-I**), suggesting that the reduction of the two proteins seen in stiffer materials with the Wnt inhibitors may be downstream of either non-canonical Wnt or other associated pathways. The siCTNNB1-associated elevation of p-Smad1/5 expression on stiffer MC was associated with an increase in BMP4 expression which was not present on NX-MC (**Figure 7J-L**). BMP2 and BMP7 expression did not appear to be affected by siCTNNB1 when compared to siControl on either scaffold, although BMP2 expression was generally higher on NX-MC and BMP7 expression was slightly higher on MC in the presence of siCTNNB1. Taken together, our data suggests that β-catenin serves to negatively regulate osteogenic differentiation, Smad1/5 phosphorylation, and BMP4 expression specifically in stiffer MC-GAG materials versus structurally and compositionally identical, but mechanically softer NX-MC scaffolds.

## Discussion

In this work, we evaluated the contributions of the canonical Wnt signaling pathway to osteogenic differentiation induced by differential stiffness of a synthetic nanoparticulate mineralized collagen glycosaminoglycan material (*3*). Using two small molecule inhibitors that, respectively, stabilized the β-catenin destruction complex (IWR1) or reduced the production of Wnt ligands (IWP2), we demonstrated that Wnt inhibition differentially affected the stiffer MC-GAG material such that the expression of activated β-catenin, osteogenic gene expression, and mineralization was reduced in a manner not observed on less stiff, non-crosslinked NX-MC scaffolds. Wnt inhibition also induced a reciprocal increase in p-Smad1/5, the canonical BMPR intracellular mediator, and Runx2 proteins preferentially on the stiffer MC-GAG materials. The increase in p- Smad1/5 is likely secondary to an increase in expression of the BMP4 ligand, which is found to occur only on stiff MC-GAG in response to IWR1. Significant interactions exist between cWnt and YAP/TAZ such that both are upregulated on the stiffer material and both are downregulated with Wnt inhibition. In addition, near complete colocalization occurs between activated β-catenin and YAP with both primarily cytosolic on soft MC-GAG and both cytosolic and nuclear on stiff MC-GAG. However, Wnt inhibition reduced the colocalization, suggesting that the activities of YAP may be separated from cWnt. Unlike treatment with the small molecule Wnt inhibitors which indirectly affect cWnt and introduce a certain amount of non-specificity, direct inhibition via β-catenin knockdown demonstrated minimal differences in the expression of the mechanotransduction mediators YAP and TAZ with a mild increase of p-FAK on the stiffer material, while all three were increased on the softer MC-GAG material. β-catenin knockdown also increased osteogenic gene expression and mineralization specifically on the stiffer material with no effects on the softer NX-MC material. While there are overall differences in Wnt inhibition versus β-catenin knockdown, one consistent effect was the increase in p-Smad1/5, Runx2, and BMP4 expression again only on the stiffer material. Our data suggests several conclusions: 1. Stiffness of MC-GAG triggered activation of the cWnt pathway, 2. Inhibition of the Wnt signaling pathways differentially affected osteogenic differentiation on stiffer MC-GAG materials, 3. While the YAP-induced mechanosensitive pathways are both influenced by stiffness and frequently parallel cWnt, separation of the two pathways can occur, 4. β-catenin functions as a negative regulator for osteogenesis in response to stiffness via downregulation of the BMPR signaling pathways and BMP4 expression.

ECM-inspired regenerative materials have gained significant attention due to the known effects of the ECM in directing progenitor cell differentiation and cell fate determination. One of the primary advantages of synthetic materials over decellularized ECM is tunability such that material properties, such as stiffness, as well as biochemical compositions may be altered to affect specific cell behaviors. However, despite such advantages, mechanistic changes in cell biological responses to specific material alterations are not well understood for several reasons. First, mechanisms differ significantly between different materials even when the compositions are similar. For example, we previously reported that osteogenic differentiation on non-mineralized versus mineralized collagen glycosaminoglycan materials fabricated in the same manner with the same organic compositions differed such that the latter had essentially no dependence on ERK1/2 pathway while the former required ERK1/2 for differentiation (*5*). Similarly, in this report we demonstrated that differential stiffness of the same material elicited a vast difference in the contribution of the Wnt pathways such that the softer materials differentiated without reliance on Wnt, albeit less efficiently compared to hMSCs within the stiffer MC scaffold. Second, differential cell types may correspond to different mechanisms thereby causing contradictory findings. There is little doubt that, in total, Wnt pathways are important in the osteogenic differentiation process induced by stiff ECM or materials as we and others have shown that overall inhibition of the pathways corresponded to reductions in osteogenesis (*29*). However, cWnt and β- catenin may be either positive or negative regulators depending upon the cell type, species, and environment for which it is studied. Several conditional knockout murine models have indicated that β-catenin is essential for osteoblast differentiation and maturation in development (*18, 30*). In contrast, there are a number of instances where others have reported that β-catenin inhibits osteogenic differentiation. Both De Boer et al and Boland et al reported that Wnt3A, the classic cWnt ligand, downregulated osteogenic differentiation of bone marrow-derived primary hMSCs (*20, 21*).

Furthermore, the latter group demonstrated that a dominant negative form of TCF1, the major transcription factor in the cWnt pathway, activated osteogenic gene expression (*20*). In the setting of periodontitis, Liu et al noted that periodontal ligament stem cells (PDLSCs) expressed higher levels of β-catenin compared to normal PDLSCs, yet displayed a decreased ability to undergo osteogenic differentiation in a manner that could be rescued by Dikkopf-1, an inhibitor of cWnt (*31*). The example of PDLSCs in periodontitis is particularly interesting as it is clearly secondary to environmental influences, an analogous situation to our current work, such that the local microenvironment has dictated the quantity of β-catenin and its role in differentiation.

The complexity of cell biological changes induced by materials is further compounded by the complicated interactions between the involved signaling pathways. With respect to the interactions between Wnt and BMP signaling, both synergistic and antagonistic effects have been reported. Mbalaviele and colleagues demonstrated that a constitutive active β-catenin mutant synergized with BMP-2 to induce *in vitro* mineralization of osteoblast cell lines and *in vivo* calvarial bone formation (*32*). Similarly, Azevedo et al reported that the decrease in BMP4 expression in hMSCs from patients with acute myeloid leukemia was associated with the decrease in β-catenin (*33*). Yet, cWnt signaling has also been established to inhibit BMP4 expression as shown in Xenopus development and murine corneal epithelium (*34, 35*). The addition of YAP/TAZ further complicates the picture. Stiffness has been well-associated with an increase in osteogenic differentiation in a manner that requires YAP/TAZ expression (*36*). Yet, YAP paradoxically has been reported to suppress Smad1/5 phosphorylation in a murine MSC cell line (*37*). The complexity of the interactions between the pathways reiterates that cell biological responses to even a single material property, such as targeted changes in stiffness, are not universal and depend upon the totality of factors combined.

The identification of β-catenin and cWnt as a negative regulator of osteogenic differentiation on stiffer MC-GAG suggests that it may be a useful molecular marker for material refinement. While stiffness is generally considered to be positively associated with osteogenic differentiation, there is likely an endpoint in which further increases in stiffness does not correlate to more osteogenic differentiation due to the expression of β-catenin. Further, the multi-scale mechanical properties of porous biomaterials suggest an exciting avenue for future analysis of cell response. While the reported stiffness of many regenerative biomaterials is a function of the material porosity, the local mechanical environment experienced by an individual cell within the material is a function of bulk stiffness as well as pore architecture (*24, 38, 39*). Here, targeted manipulation of structural and mechanical parameters in architecture may yield a material with optional macro-scale mechanical performance to meet surgical practicality needs, while also unlocking a new design space to modulate cell response in the context of local, micro-scale mechanical properties. Clinically, this realization is likely beneficial as implantation of excessively-stiff materials for bone regeneration is not particularly desirable due to the potential for fracture, inability to conform to variable defects, and potential limitations of tissue ingrowth.

## Materials and Methods

### Fabrication of Mineralized Collagen Scaffolds

MC-GAG scaffolds were prepared using the lyophilization process described previously (*23–25*). Briefly, a suspension of collagen and GAGs were produced by combining microfibrillar, type I collagen (Collagen Matrix, Oakland, NJ) and chondroitin-6-sulfate (Sigma-Aldrich, St. Louis, MO) with calcium salts (calcium nitrate hydrate: Ca(NO3)2·4H2O; calcium hydroxide: Ca(OH)2, Sigma-Aldrich, St. Louis, MO) in a solution of phosphoric acid. The suspension was frozen using a constant cooling rate technique (1 °C/min) from room temperature to a final freezing temperature of −10 °C using a freeze dryer (Genesis, VirTis). Following sublimation of the ice phase, scaffolds were sterilized via ethylene oxide and cut into 6 or 8 mm diameter disks for culture. NX- MC scaffolds were rehydrated with phosphate buffered saline (PBS) overnight and used for cell culture. For stiffer MC scaffolds, scaffolds were rehydrated in PBS overnight with 1-ethyl-3-(3- dimethylaminopropyl) carbodiimide (EDAC, Sigma-Aldrich) and N-hydroxysuccinimide (NHS, Sigma Aldrich) at a molar ratio of 5:2:1 EDC:NHS:COOH where COOH represents the amount of collagen in the scaffold as we previously described (*26*).

### Cell Culture

Primary hMSCs (Lonza, Inc., Allendale, NJ) were expanded in Dulbecco’s Modified Eagle’s medium (DMEM, Corning Cellgro, Manassas, VT) supplemented with 10% fetal bovine serum (Atlanta Biologicals, Atlanta, GA), 2 mM L-glutamine (Life Technologies, Carlsbad, CA), 100 IU/mL penicillin/100 μg/mL streptomycin (Life Technologies). After expansion, 3 x 10^5^ hMSCs were seeded onto 8 mm NX-MC and MC scaffolds in growth media. 24 h after seeding, media was switched to osteogenic differentiation media consisting of 10 mM β-glycerophosphate, 50 μg/mL ascorbic acid, and 0.1 μM dexamethasone (Sigma Aldrich, St. Louis MO). Cells were treated with the Wnt antagonists IWR1-endo or IWP2 (EMD Millipore Corp, Burlington, MA) at the specified concentrations for 3 days in the two-dimensional cultures or 7 days, 14 days, or 8 weeks on NX-MC or MC.

### Quantitative Real-time Reverse-Transcriptase Polymerase Chain Reaction

Total RNA was extracted using the RNeasy kit (Qiagen, Valencia, CA) at 0, 3, and 7 days of culture. Gene sequences were obtained from the National Center for Biotechnology Information gene database and primers were designed (Eurofins Genomics, Louisville, KY, Supplemental Table 1). Quantitative real time RT-PCR (QPCR) was performed on the Opticon Continuous Fluorescence System (Bio-Rad Laboratories, Inc., Hercules, CA) using the QuantiTect SYBR Green RT-PCR kit (Qiagen). Cycle conditions were as follows: reverse transcription at 50°C (30 min); activation of HotStarTaq DNA polymerase/inactivation of reverse transcriptase at 95°C (15 min); and 45 cycles of 94°C for 15s, 58°C for 30 s, and 72°C for 45 s. Results were analyzed and presented as representative graphs of triplicate experiments.

### Western Blot

Cultured cells at 0 and 3 days of culture or scaffolds at 0 and 7 days of culture were resuspended in 3x SDS reducing sample buffer to create total protein lysates. Lysates were incubated at 95°C for 5 minutes and then centrifuged in 0.2 μm Spin-X filters (Corning, Costar, Corning, NY) at 14,000 rpm for 5 minutes. Protein concentrations were measured and subjected to SDS-polyacrylamide gel electrophoresis (Bio-Rad, Hercules, CA) at 4 to 20%. Western blot analysis was performed with antibodies against, phosphorylated Smad1/5 (p-Smad1/5), total Smad5, phosphorylated ERK1/2 (p-ERK1/2), total ERK1/2, phosphorylated Akt (p-Akt), total Akt, phosphorylated FAK (p- FAK), total FAK, phosphorylated p38 (p-p38), total p38, Runx2, YAP, TAZ, non-p-β- catenin, glyceraldehyde-3-phosphate dehydrogenase (GAPDH), and β-actin, followed by 1:4000 dilutions of HRP-conjugated immunoglobulin G antibodies (Bio-Rad, Hercules, CA). Detection was performed using an enhanced chemiluminescent substrate (Thermo Scientific, Rockford, IL). All primary antibodies against phosphorylated proteins were obtained from Cell Signaling Technologies (Beverly, MA), and all primary full-length antibodies were obtained from Santa Cruz Biotechnology (Santa Cruz, CA). Imaging was carried out and quantified using Image J (NIH, Bethesda, MD).

### Microcomputed Tomography

Scaffolds were fixed using 10% formalin and mineralization was quantified by micro-computed tomographic imaging (microCT) using Scanco 35 (Scanco Medical AG, Bruttisellen, Switzerland) in triplicate. Scans were performed at medium resolution with a source voltage of 70 E (kVp) and I (μA) of 114. The images had a final element size of 12.5 μm. Images were analyzed using software supplied from Scanco (Image Processing Language version 5.6) and reconstructed into three-dimensional (3D) volumes of interest. Optimum arbitrary threshold values of 20 (containing scaffold and mineralization) and 80 (containing mineralization alone) were used uniformly for all specimens to quantify mineralized areas from surrounding unmineralized scaffold. Analysis of 3D reconstructions was performed using Scanco Evaluation script #2 (3D segmentation of two volumes of interest: solid dense in transparent low-density object) and script #6 (bone volume/density only bone evaluation) for volume determinations.

### Alizarin Red Staining

Scaffolds were fixed in 10% formalin, paraffin-embedded, and sectioned at 4 microns. Following deparaffinization, sections were stained with 1 mL of 1 mg/mL of Alizarin Red Stain solution (Acros Organics, Fair Lawn, NJ) for 30 minutes. Images were captured with the Zeiss Axio Observer 3 inverted microscope with the ZEN 2.3 Pro software (Zeiss, Oberkochen, Germany). For quantitative analysis, stained slides were incubated with 500 μL of 10% glacial acetic acid (ThermoFisher, Waltham, MA) for 30 minutes. The samples were centrifuged and neutralized with 10% ammonium hydroxide to a pH between 4.1-4.5. Absorbance was measured at OD405 (Epoch spectrophotometer, BioTek, Winooski, VT).

### Confocal Microscopy

Scaffolds were fixed in 10% formalin, paraffin-embedded, and sectioned at 4 μm. Following deparaffinization, the sections were treated with 0.5% Triton X-100 (MP Biomedicals, LLC, Santa Ana, CA), blocked with 10% normal goat serum (Jackson ImmunoResearch Laboratories, Inc, West Grove, PA) for 1 hour, and subjected to heat-induced antigen retrieval using sodium citrate buffer(10 mM, pH 6, ThermoFisher, Waltham, MA) at >80°C, for 20 minutes. Sections were then separately incubated with anti-Yap (1:200, Santa Cruz Biotechnology, Santa Cruz, CA) anti-non-p-β-catenin (1:1000 Cell Signaling Technologies, Beverly, MA) overnight at 4 °C. After washing, sections were incubated in anti-rabbit IgG Alex Fluor Plus 488 or anti-mouse IgG Alexa Fluor Plus 594 (ThermoFisher, Waltham, MA). Coverslips were mounted with Prolong Gold Antifade Reagent with Dapi (Cell Signaling Technologies, Danvers, MA). Images were captured with the Zeis LSM900 confocal laser scanning microscope with Zen 3.1 Blue software (Zeiss, Oberkochen, Germany).

### RNA Interference

2.5 x 10^4^ hMSCs were seeded per well in 24 well plates for 24 hours and transfected with Lipofectamine RNAiMAX (ThermoFisher, Waltham, MA) using the manufacturer’s protocol. Briefly, 1.5 μL of reagent and 50 pmol of scrambled small interfering RNA (siRNA) for control (siControl, Santa Cruz Biotechnology, Santa Cruz, CA) or siRNA targeting CTNNB1 (siCTNNB1, targeting sequence 5′- CCACAGCUCCUUCUCUGAGUGGUAA -3’, ThermoFisher, Waltham, MA) were resuspended in 50 μL of Opti-MEM and incubated at room temperature for 30 min. Each well was then incubated with the Lipofectamine/siRNA solution for 5 hours, after which 1 mL of DMEM with 10% FBS was added. 24 hours after transfection, cells were harvested and 1 x 10^5^ of transfected hMSCs were seeded onto 6 mm NX-MC and MC scaffolds.

### Statistical Analysis

Statistical analyses were performed using SPSS Version 27 (Chicago, IL) using independent samples Student’s t test or analyses of variance (ANOVA) followed by post hoc tests under the Tukey criterion for samples with equal variances. For multiple comparisons with samples of unequal variances, Welch’s ANOVA with post hoc tests under the Games-Howell criterion were performed (Supplemental Table 2). A value of p<0.05 was considered significant.

## Funding

This work was supported by the National Institutes of Health/National Institute of Dental and Craniofacial Research under award numbers R01 DE028098 (JCL), R01 DE029234 (JCL), R21 DE026582 (BACH), the Bernard G. Sarnat Endowment for Craniofacial Biology (JCL), and the Jean Perkins Foundation (JCL). The content is solely the responsibility of the authors and does not necessarily represent the official views of the National Institutes of Health.

## Author contributions

JCL designed the experiments and interpreted the data. QZ, XR, MKO performed the experiments and analyzed of the data. MJD, VK, and BACH fabricated the scaffolds. BACH contributed valuable comments at multiple stages of the study. JCL, QZ, MKO, and RMC wrote the manuscript and all authors contributed to review of the manuscript.

## Competing Interests

The authors have no financial or competing interests to disclose.

## Data and materials

All data needed to evaluate the conclusions in the paper are present in the paper and/or the Supplementary Materials. Additional data available from authors upon request.

## Supplementary Materials

**Supplemental Table 1.**
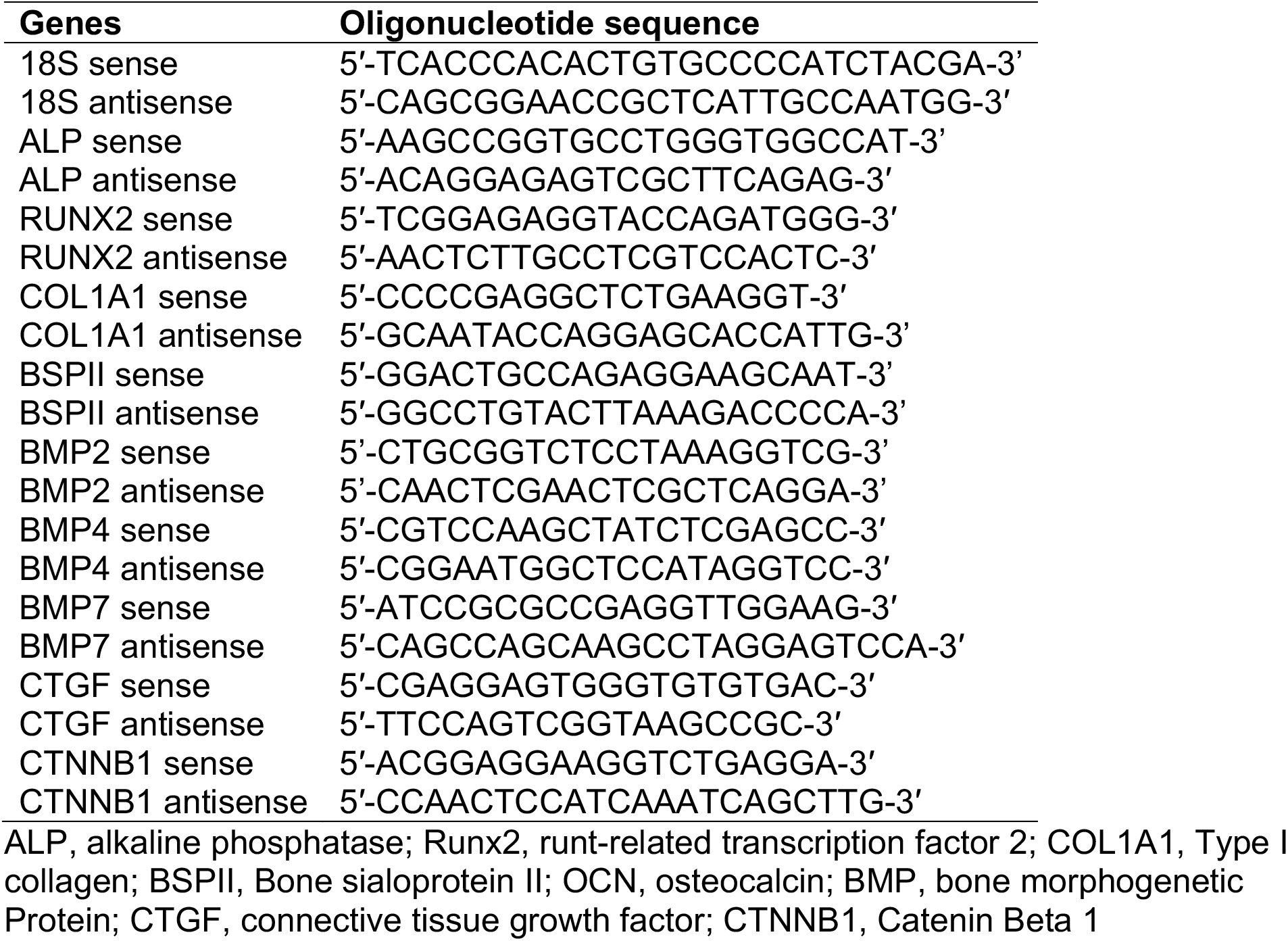
Primer Sequences.

**Supplemental Table 2.**
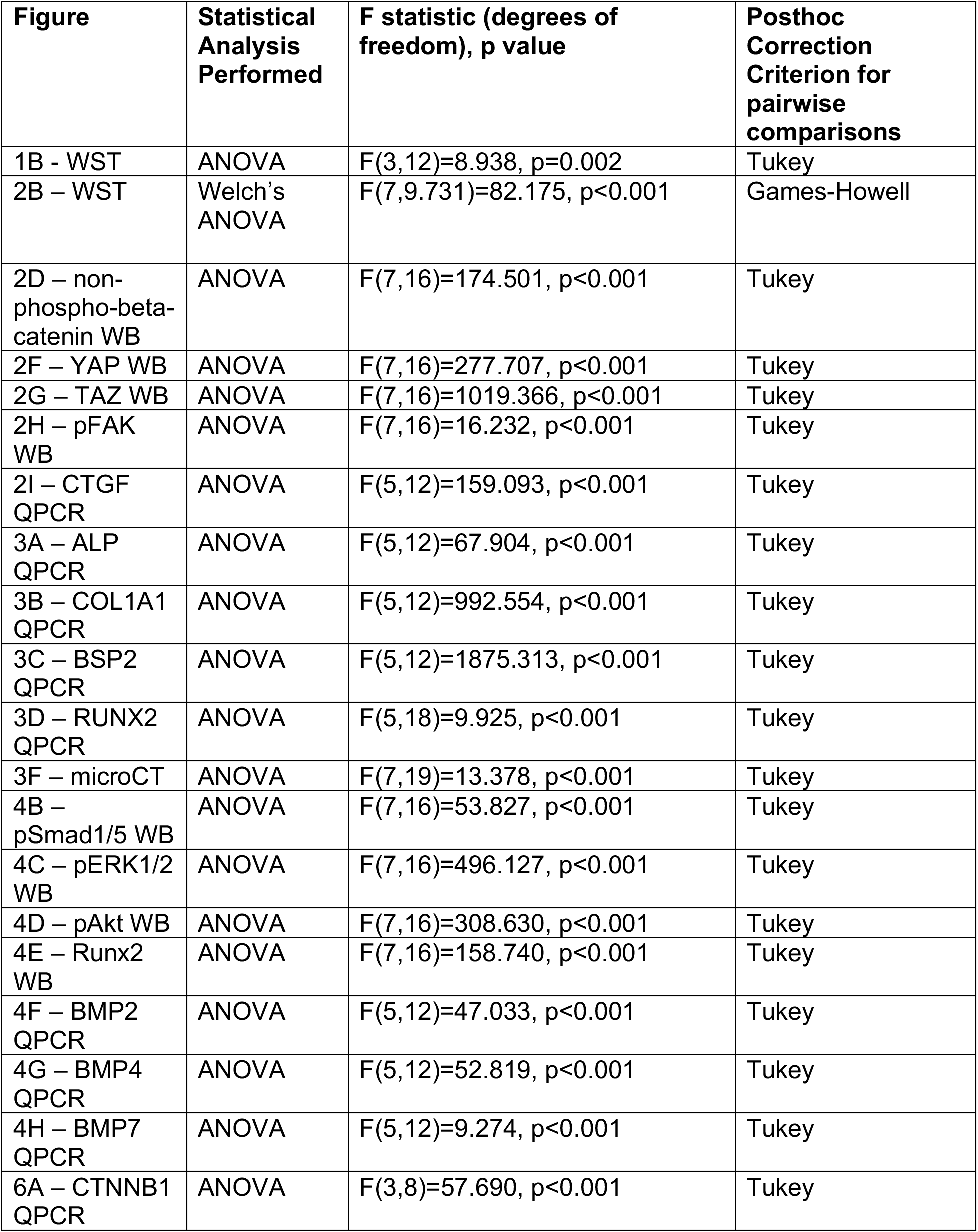
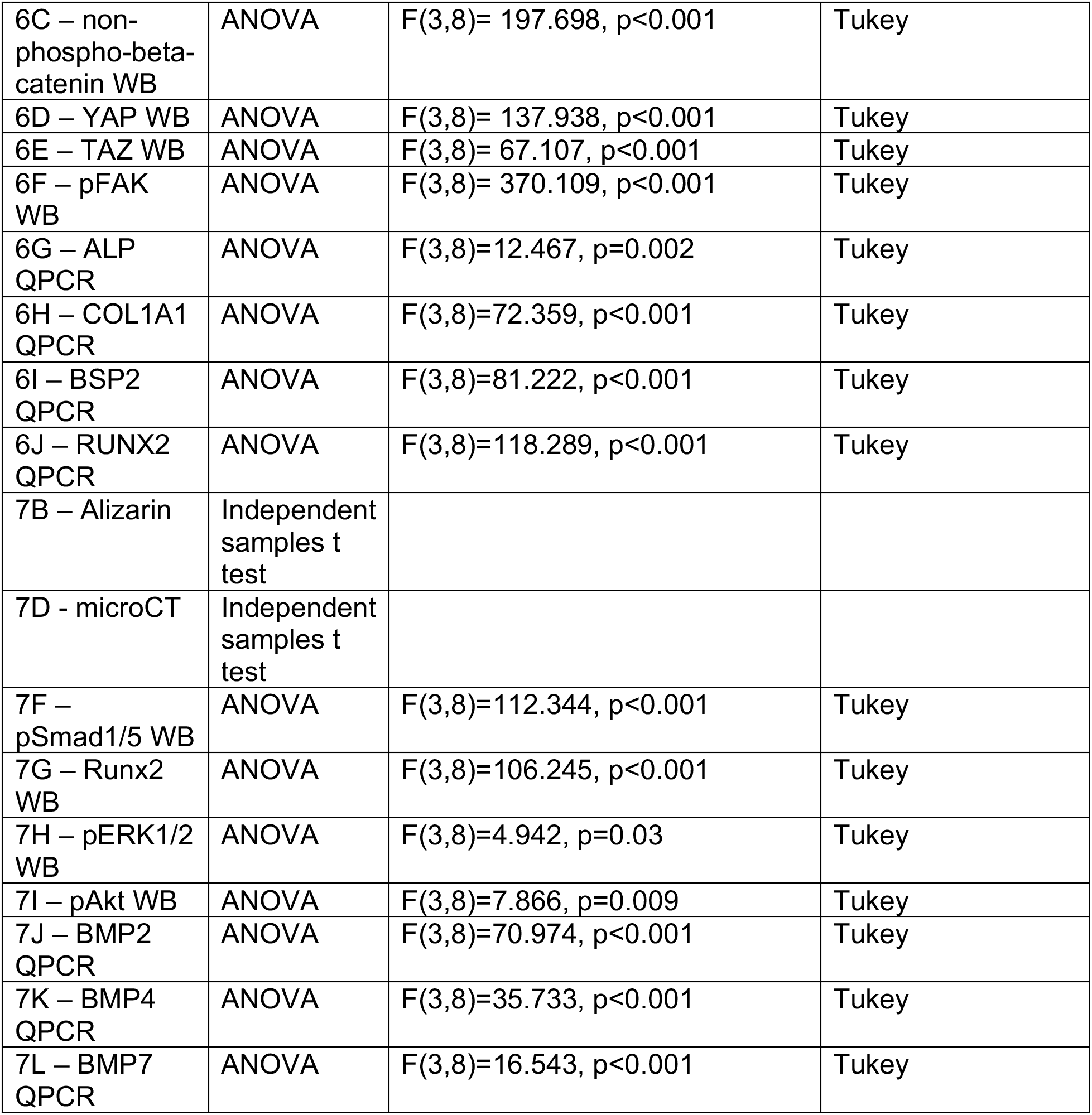
Summary of Statistical Analyses.

## Notes

### Competing Interest Statement

The authors have declared no competing interest.

